# *Plasmodium falciparum* sexual parasites regulate infected erythrocyte permeability

**DOI:** 10.1101/2020.05.18.101576

**Authors:** Guillaume Bouyer, Daniela Barbieri, Florian Dupuy, Anthony Marteau, Abdoulaye Sissoko, Marie-Esther N’Dri, Gaelle Neveu, Laurianne Bedault, Nabiha Khodabux, Diana Roman, Sandrine Houzé, Giulia Siciliano, Pietro Alano, Rafael M. Martins, Jose-Juan Lopez-Rubio, Jérome Clain, Romain Duval, Stéphane Egée, Catherine Lavazec

**Affiliations:** Sorbonne Université, CNRS UMR 8227, Station Biologique de Roscoff, France; Laboratoire d’excellence GR-Ex, France; Université de Paris, Inserm U1016, CNRS UMR 8104, Institut Cochin, France; Université de Paris, IRD 261, MERIT, France; Istituto Superiore di Sanita, Roma, Italy; Université de Montpellier 1 & 2, CNRS 5290, IRD 224, MIVEGEC, France

**Author notes:** contributed equally to this work (Guillaume Bouyer, Daniela Barbieri). contributed equally to this work (Stéphane Egée, Catherine Lavazec).

## Abstract

To ensure the transport of nutrients necessary for their survival, *Plasmodium falciparum* parasites increase erythrocyte permeability to diverse solutes. These New Permeation Pathways (NPP) have been extensively characterized in the pathogenic asexual parasite stages, however the existence of NPP has never been investigated in gametocytes, the sexual stages responsible for transmission to mosquitoes. Here, we show that NPP are still active in erythrocytes infected with immature gametocytes and that this activity declines along gametocyte maturation. Our results indicate that NPP are regulated by cyclic AMP (cAMP) signaling cascade during sexual parasite stages, and that the decrease in cAMP levels in mature stages results in a slowdown of NPP activity. We also show that NPP facilitate the uptake of artemisinin derivatives and that phosphodiesterase (PDE) inhibitors can reactivate NPP and increase drug uptake in mature gametocyte-infected erythrocytes. These processes are predicted to play a key role in *P. falciparum* gametocyte biology and susceptibility to antimalarials.

## INTRODUCTION

Malaria remains a major public health problem with more than 200 million cases and almost half a million deaths annually. The clinical symptoms of malaria are attributed to *Plasmodium* asexual stages, whereas parasite transmission from humans to mosquitoes relies on the gametocytes, the specialized sexual cells formed by a fraction of parasites which cease asexual propagation. *Plasmodium falciparum* gametocyte maturation requires about ten days, and is classically divided into five developmental stages based upon morphological features (*1*). If the fight against malaria obviously requires compounds targeting asexual stages, transmission remains the Achilles heel of the strategies implemented today and are a key target in WHO agenda towards malaria eradication (*2*). Most antimalarial drugs, including artemisinin derivatives, target the asexual stages but are less effective against gametocytes (*3, 4*). As a consequence, infected individuals remain a source of transmission even after they are cured. It remains unclear why gametocytes become less sensitive to artemisinin derivatives as their maturation progresses from stage I to stage V. Thus, understanding the biology of gametocyte development within erythrocytes is crucial for successful malaria elimination.

Following *P. falciparum* invasion, the infected erythrocyte displays important alterations of its membrane properties. For instance, asexual parasites increase erythrocyte permeability to diverse solutes to ensure the transport of nutrients and waste products necessary for their replication and survival. Mature asexual parasites activate weakly selective anion channels in the erythrocyte membrane to generate New Permeability Pathways (NPP) that render infected erythrocytes more permeable to a range of nutrients (*5, 6*) and to several antimalarials (*7–10*). Key proteins involved in NPP have been characterized, like parasitic proteins CLAG3/RhopH1, RhopH2 and RhopH3 or endogenous constituent of the peripheral-type benzodiazepine receptor (PBR) (*11–14*). However, the full identity of the NPP has never been conclusively established (*15*). Although their regulatory mechanisms also remain unclear, cyclic AMP (cAMP)/Protein Kinase A (PKA) pathway seems to play a key part in regulating ion channels in the membrane of erythrocytes infected with asexual stages (*16*).

Despite the vital role of NPP in asexual stages, NPP level of activity in gametocyte-infected erythrocytes (GIE) is unknown, nor is the role of NPP in antimalarials uptake by GIE. Refractoriness of GIE to lysis upon short exposition to isosmotic sorbitol solution has led to the dogma that NPP are totally absent in gametocyte stages (*17*). However, this assumption is not supported by the fact that during their ten-day maturation, gametocytes also need to absorb nutrients from the plasma and get rid of toxic waste products they generate upon hemoglobin digestion, two major roles of NPP during asexual stages. In addition, absence of NPP in GIE is not consistent with the expression profile of members of the RhopH complex that are synthesized in mature intracellular parasites and then secreted upon egress onto the erythrocyte targeted for invasion (*18, 19*). Therefore, all known NPP components should be present at the surface of newly invaded GIE.

In this study, we performed isosmotic lysis, electrophysiology, fluorescence tracer uptake and viability experiments to evaluate NPP activity during *P. falciparum* gametocytogenesis. We found that NPP activity is regulated by cyclic AMP signaling cascade, and interfering with this pathway can reactivate erythrocyte permeability and facilitate uptake of artemisinin derivatives by mature gametocytes.

## RESULTS

### NPP are still active in immature gametocytes

To evaluate the NPP activity in GIE, we first performed measurements of isosmotic lysis of immature GIE in sorbitol, a sugar alcohol permeant through NPP (*6*). Sorbitol uptake was drastically reduced in stage II GIE compared to trophozoites and schizonts, however about 36% GIE were lysed after 60 minutes in sorbitol, suggesting that erythrocyte permeability is modified by immature gametocytes (Fig. 1a). Lysis kinetics in stage II GIE were similar to that of late rings 16 hours-post-invasion, when NPP start to be active during the asexual cycle (*20*). GIE lysis was significantly inhibited by the general anion channel inhibitors 5-nitro-2-(3-phenylpropylamino)benzoic acid (NPPB) and furosemide, by the benzodiazepine Ro5-4864 and by the isoquinoline PK11195, which have all been shown to inhibit NPP (Fig. 1b) (*11, 21*). Equivalent lysis kinetics observed in alanine or phenyltrimethylammonium (PhTMA^+^) isosmotic solutions, with a slightly greater lysis in PhTMA^+^ than in sorbitol, confirmed that uptake occurred through NPP (Fig. c). Measurement of membrane currents using whole-cell patch-clamp strengthened observations obtained with isosmotic lysis. Quantification of total ion fluxes across the host membrane of individual infected erythrocytes indicated that membrane conductance at −100 mV was 6-fold lower in stage II GIE than in trophozoites (Fig. 1d). However, conductance in early GIE showed anion selectivity, inward rectification and NPPB sensitivity (Fig. 1e), which are three fundamental characteristics of the correlate membrane currents of NPP activity (*22*). Pharmacological inhibition of NPP is expected to lead to nutrient starvation and accumulation of toxic metabolic wastes eventually leading to parasite death. Thus, we measured viability of early gametocytes (stages I-II) upon NPP inhibition by using a luciferase-based assay with a transgenic parasite expressing luciferase under the control of the gametocyte-specific promoter *Pfs16* (*23*). We observed a slight but significant decrease in early gametocytes viability 48 hours after a 3 hours exposure to 100 µM NPPB, consistent with a vital role of NPP activity (Fig. 1f). Therefore, organic solute uptake, patch-clamp and viability studies indicate that NPP are weaker than in trophozoites but are still active and necessary in early GIE.

**Figure 1.**
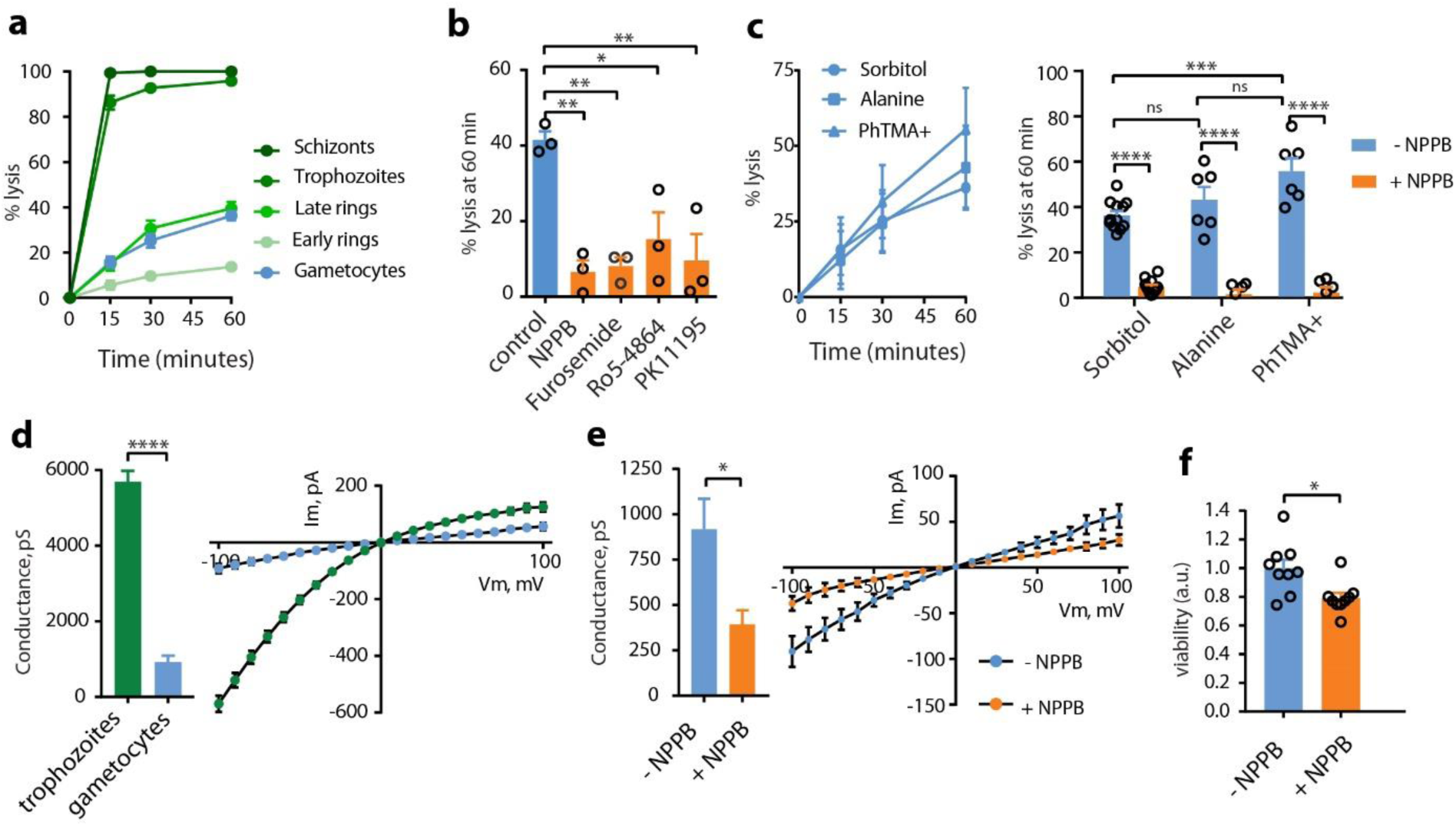
NPP are still active in immature gametocytes. **a.** Kinetics of sorbitol-induced isosmotic lysis of erythrocytes infected with early rings (4 hours-post-invasion (hpi)), late rings (16 hpi), trophozoites (28hpi), schizonts (40hpi) and stage II gametocytes during 60 minutes. n = 3 experiments. **b.** % lysis of stage II GIE in sorbitol at 60 minutes with or without 100 µM NPPB, Furosemide, Ro5-4864, or PK11195. **c.** Lysis kinetics (left) and % lysis at 60 minutes (right) of stage II GIE in sorbitol, alanine or PhTMA^+^. **d.** Patch clamp experiments on erythrocytes infected with trophozoites or stage II gametocytes (number of cells = 14 and 14, respectively). Left: whole cell conductance calculated at −100 mV. Right: I-V plot from patch clamp experiments. **e.** Patch clamp experiments on stage II GIE in presence or absence of NPPB (number of cells = 14 and 5, respectively). Left: whole cell conductance calculated at −100 mV. Right: I-V plot from patch clamp experiments. **f.** Viability (luciferase activity) of early gametocytes of the NF54-cg6-pfs16-CBG99 line 48 hours after a 3-hour incubation with or without 100 µM NPPB. The graph shows relative viability normalized by the average luciferase activity of control (without NPPB). a.u.: arbitrary units. In b, c, f, circles indicate the number of independent experiments. Error bars show the standard error of the mean (SEM). Statistical significance is determined by a Mann Whitney test (d, e, f) or by one-way ANOVA with Dunnet correction (b) or Sidak correction (c) for multiple comparisons.

### NPP decline along gametocytogenesis

To further analyze the evolution of this permeability along gametocytogenesis, we performed isosmotic lysis experiments in sorbitol, alanine and PhTMA^+^ for synchronous cultures of stage I, II, III, IV, and V gametocytes (Fig. 2a-c). A transgenic parasite expressing GFP under the control of the *Pfs16* promoter was used to discriminate stage I gametocytes from asexual stages (*24*). From stage I to stage V, we observed a progressive decrease in permeability for the three different solutes, leading to undetectable NPP activity in mature GIE. Electrophysiological experiments using whole cell configuration performed on GIE from all stages confirmed that membrane conductance drops along gametocyte maturation, leading to currents in mature GIE corresponding to levels usually recorded in uninfected erythrocytes (Fig. 2d and Supp. Fig. S1). Altogether, these data indicate that NPP activity declines along gametocytogenesis.

**Figure 2.**
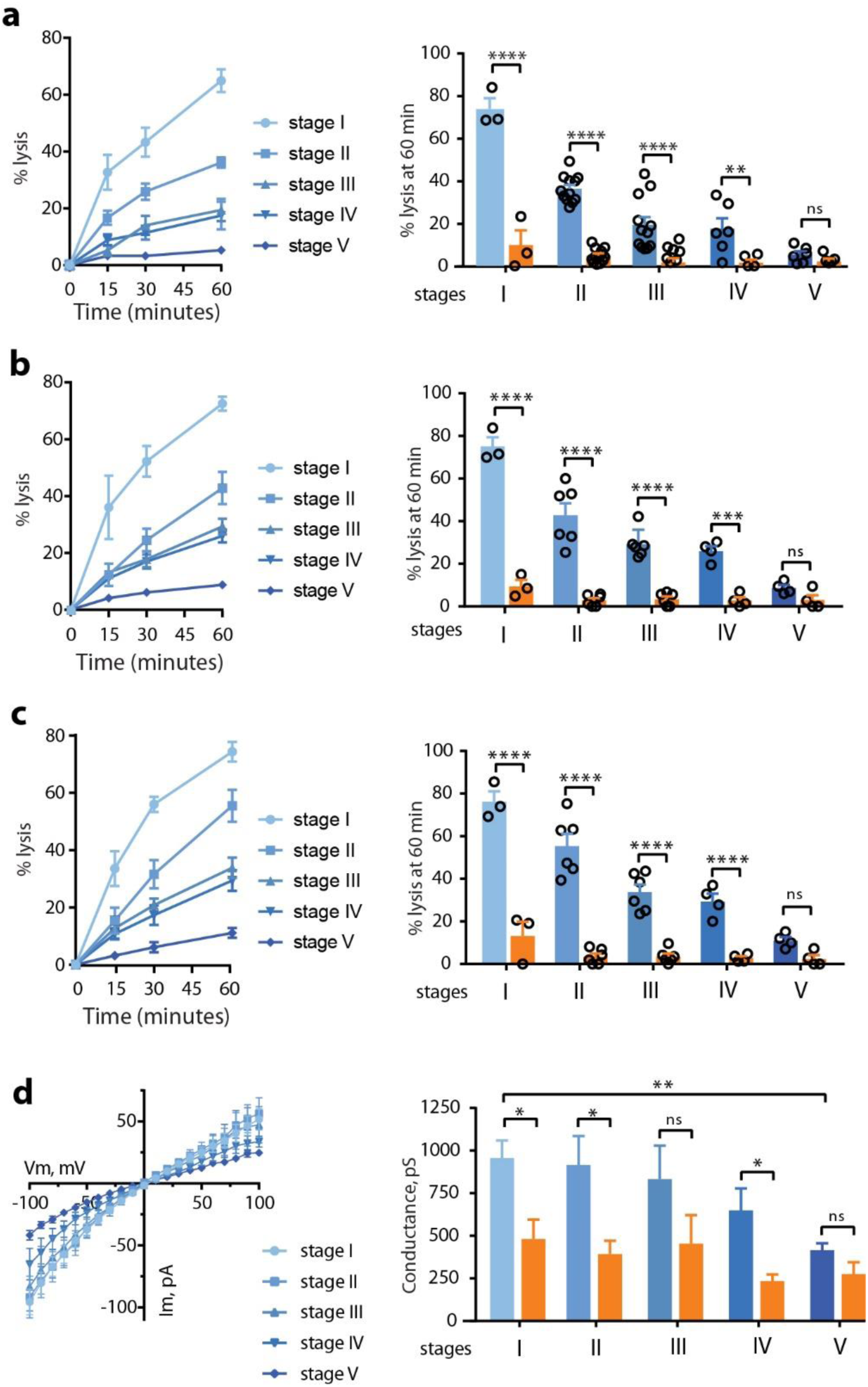
NPP decline along gametocytogenesis. **a-c.** Sorbitol-induced (a), Alanine-induced (b) and PhTMA^+^-induced (c) isosmotic lysis of GIE from stage I to stage V. Left: Kinetics of isosmotic lysis during 60 minutes. Right: % lysis at 60 minutes with (orange) or without (blue) 100 µM NPPB. Circles indicate the number of independent experiments and error bars show the SEM. Statistical significance is determined by one-way ANOVA with Sidak correction for multiple comparisons. **d.** Left: I-V plot from patch experiments on GIE from stage I to stage V. Right: Whole cell conductance calculated at −100 mV on GIE from stage I to stage V with (orange) or without (blue) 100 μM NPPB. Error bars show the SEM. Statistical significance is determined by a Mann Whitney test.

### NPP activity is regulated by cAMP-signaling

We then aimed to determine the mechanisms underlying the regulation of NPP activity in GIE. We have previously observed that cAMP/PKA pathway activates ion channels in the membrane of erythrocytes infected with asexual stages (*16*), suggesting that cAMP-signaling may also modulate NPP during gametocytogenesis. To address this hypothesis, we performed sorbitol lysis experiments on early GIE pre-incubated with KT5720 and H89, two independent and widely used PKA inhibitors that have already been shown to inhibit PKA activity in *P. falciparum* (*25, 26*). We found that both compounds strongly ablated GIE permeability (Fig. 3a), whereas the cAMP analogue 8-bromoadenosine-3’-5’-cyclic monophosphate (8Br-cAMP) significantly increased the sorbitol-induced lysis (Fig. 3b). As an alternative way to investigate the effects of inhibiting PKA activity, we used a transgenic parasite that overexpresses the regulatory subunit of PKA using an episome selected by the antimalarial drug pyrimethamine (pHL*pfpkar*) (*16, 26*). The down-regulation of PKA activity in this parasite line also resulted in a significant decrease in permeability of early GIE (Fig. 3c). This phenotype was reverted to the levels of wild-type parasites upon incubation with 8Br-cAMP, or in a revertant parasite line that has shed the overexpressing episome (*26*) (Fig. 3c). We observed a similar inhibition of alanine- or PhTMA^+^-induced isosmotic lysis of stage II GIE for the pHL*pfpkar* line (Supp. Fig. S2). These results indicate that PKA activity contributes to NPP in immature GIE, as described for asexual stages (*16*).

**Figure 3.**
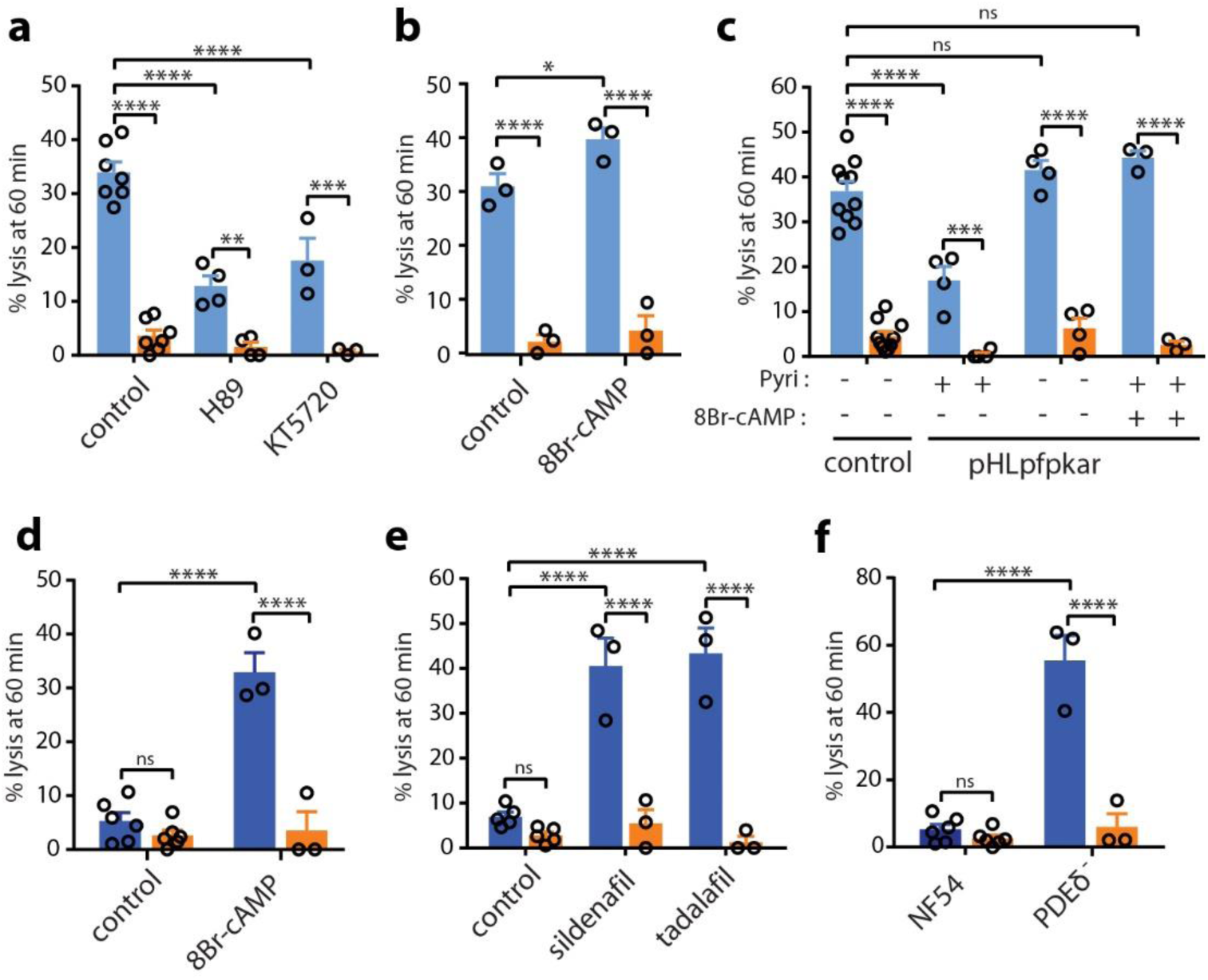
NPP activity is regulated by cAMP-signaling. **a-b.** Sorbitol-induced isosmotic lysis of stage II GIE with 100 µM H89 or 10 µM KT5720 (a), or with 100 µM 8Br-cAMP (b). **c.** Sorbitol-induced isosmotic lysis of stage II GIE from the NF54 isolate (Control) and the transgenic pHL*pfpkar* line, cultivated with or without pyrimethamine (Pyri), or pre-incubated with 100 µM 8Br-cAMP. **d-e.** Sorbitol-induced isosmotic lysis of stage V GIE with 100 µM 8Br-cAMP (d), sildenafil (e) or tadalafil (e). **f.** Sorbitol-induced isosmotic lysis of stage V gametocytes from the NF54 isolate and the transgenic line PDEδ^−^. All experiments were performed in presence (orange) or absence (blue) of 100 µM NPPB. Circles indicate the number of independent experiments and error bars show the SEM. Statistical significance is determined by one-way ANOVA with Sidak correction for multiple comparisons.

These observations suggest that the decline in NPP activity during gametocytogenesis likely results from the previously described drop in cAMP concentration in GIE due to the rising expression of PfPDEδ in mature gametocytes (*26*). Thus, we hypothesized that interfering with cAMP pathway with molecules that raise cAMP levels may turn on NPP activity in stage V GIE. As expected, raising cAMP levels in mature GIE upon incubation with 8Br-cAMP or with the PDE inhibitors sildenafil and tadalafil restored the sorbitol-induced lysis to the levels observed in early GIE (Fig. 3d, Fig. 3e and Supp. Fig. S3). Consistently, the loss of PDE activity in a transgenic parasite in which the *Pfpdeδ* gene had been deleted (*27*) also resulted in a drastic increase in permeability of mature GIE (Fig. 3f and Fig. S2). These findings show that stimulating the cAMP/PKA pathway can reactivate NPP in mature stages.

### NPP contribute to the uptake of artemisinin derivatives by immature GIE

*P. falciparum* mature gametocytes are known to become less sensitive to several antimalarials, including artemisinin derivatives, as their maturation progresses from stage I to stage V (*28, 29*). Although lower metabolic activity and complete hemoglobin digestion in mature stages have been suggested to contribute to this decrease in chemosensitivity (*4, 30*), we hypothesized that this decrease may also be linked to the decline of host membrane permeability during gametocyte maturation. To address this hypothesis, we first evaluated whether NPP activity is required for uptake of artemisinin derivatives by immature GIE. For this purpose, we used Fluo-DHA, a fluorescent probe mimicking the clinical antimalarials dihydroartemisinin (DHA) and artemether (*31*). This probe corresponds to an HPA-NBD (4-(1′-hydroxypropyl-3′-amino)-7-nitrobenzoxadiazole) spacer-fluorophore moiety attached onto the C-10 center of DHA (Supp. Fig. S4). First we validated that Fluo-DHA was detectable in the gametocyte cytoplasm after 2 hours incubation and exhibited antiplasmodial activity against early gametocytes as assessed by a luciferase-based viability assay (Supp. Fig. S4). Flow cytometry quantification showed that Fluo-DHA was embedded by about 80% of stage II GIE, whereas uninfected erythrocytes were all negative (Fig. 4a, Fig. 4b and Supp. Fig. S5). The pharmacological specificity of this probe was validated by a competition assay with an excess of DHA (Fig. 4c). Importantly, pre-incubation with the NPP inhibitors NPPB and furosemide induced a significant decrease in the proportion of positive cells, indicating that Fluo-DHA uptake is partly mediated by NPP in early GIE (Fig. 4d). About 40% cells remained fluorescent upon NPPB treatment, suggesting that Fluo-DHA uptake may also partly occur by diffusion through the erythrocyte membrane or by another route. Moreover, the viability at 48 hours of early GIE following a 3-hours pulse exposure to 150 nM Fluo-DHA or 700 nM artemisinin was slightly but significantly increased upon pre-incubation with NPPB, suggesting that NPP inhibition impacts the transport of these antimalarials into the infected erythrocytes (Fig. 4e). These results indicate that NPP facilitate the uptake of artemisinin derivatives, and substantiate the idea that the slowdown of NPPs activity in mature stages may account for their refractoriness to these drugs. To address this hypothesis, we evaluated Fluo-DHA uptake in synchronous cultures of stage II, III, IV, and V gametocytes. As observed for the decline in NPP activity, Fluo-DHA uptake also decreased during gametocyte maturation (Fig. 4f). The proportion of Fluo-DHA-positive GIE slowed down in stage III and reached the same level as that of NPPB-treated cells in stage V GIE. This profile correlates with the previously described chemosensitivity shift occurring at the transition from stage III to stage IV (*28, 32*). Therefore, the decline in NPP activity and in drug uptake along gametocytogenesis parallels the decrease in gametocytes sensitivity to several antimalarials (*4, 28*). Altogether, these results provide evidence that NPP facilitate the uptake of artemisinin derivatives in immature gametocyte-infected erythrocytes, and contribute to their chemosensitivity.

**Figure 4.**
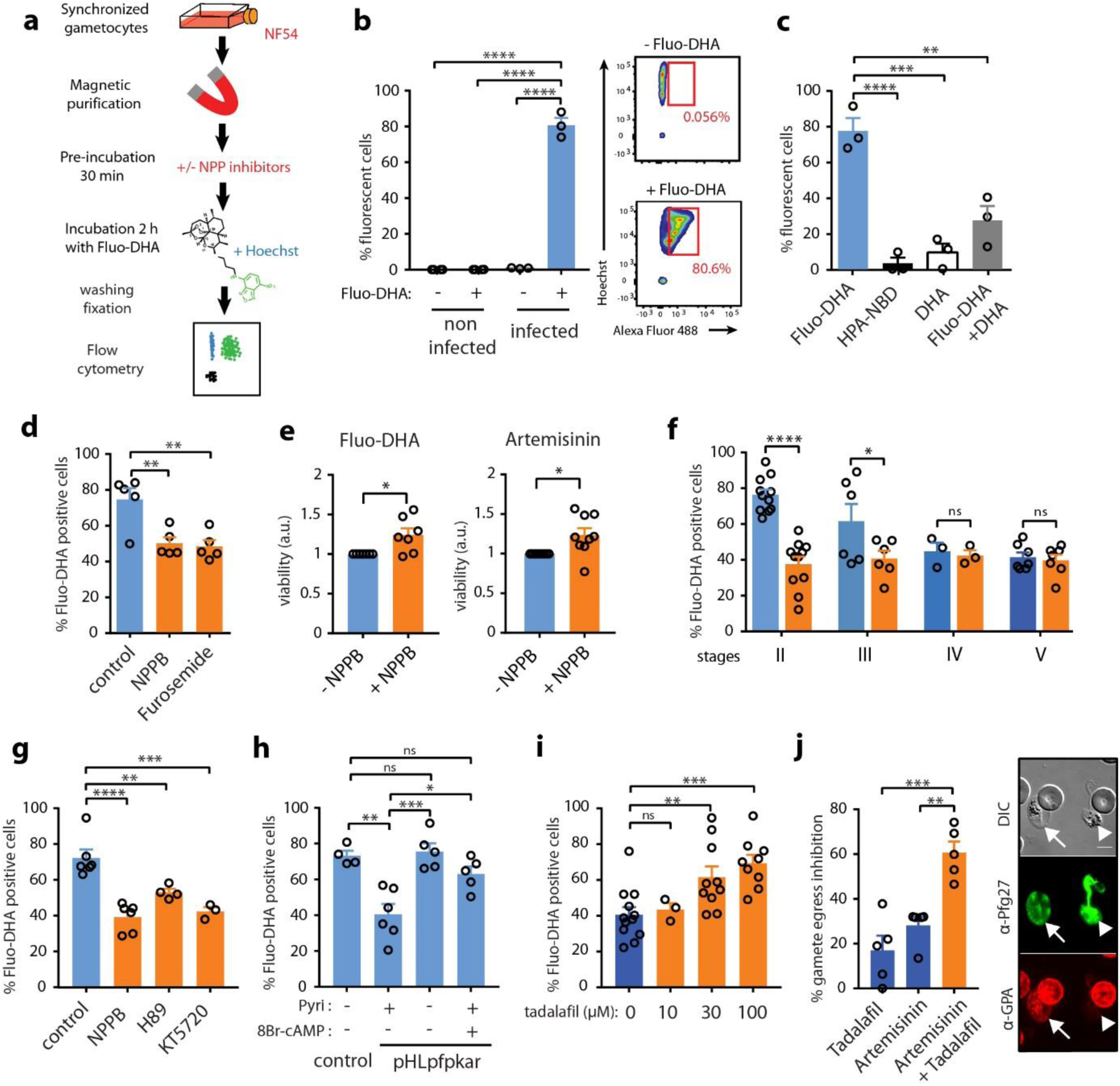
NPP contribute to the uptake of artemisinin derivatives. **a.** Diagram illustrating the uptake assay. **b.** Left: Quantification of Fluo-DHA uptake in uninfected erythrocytes and in early GIE by flow cytometry. Right: scatter plots showing the gating strategy for Fluo-DHA uptake. **c.** Competition assay between Fluo-DHA and DHA, and control of HPA-NBD and DHA-induced fluorescence levels. **d.** Inhibition of Fluo-DHA uptake in early GIE upon 100 µM NPPB or Furosemide incubation. **e.** Viability (luciferase activity) of early gametocytes of the NF54-cg6-pfs16-CBG99 line 48 hours after a 3-hour incubation with 150 nM Fluo-DHA or 700 nM artemisinin, with or without 100 µM NPPB. The graph shows the ratio of luciferase activity (drug-treated/control) and is normalized to the condition without NPPB. a.u.: arbitrary units. **f**. Quantification of Fluo-DHA uptake during gametocytogenesis with (orange) or without (blue) 100 µM NPPB. **g.** Fluo-DHA uptake in early GIE upon 100 µM H89 or 10 µM KT5720 incubation. **h**. Fluo-DHA uptake in early GIE from NF54 (Control) and the transgenic pHL*pfpkar* line, cultivated with or without pyrimethamine (Pyri), or pre-incubated with 100 µM 8Br-cAMP. **i.** Fluo-DHA uptake in stage V GIE upon 0, 10, 30 and 100 µM tadalafil incubation. **j.** Left: % inhibition of gamete egress after a 24-hour incubation with 5 µM artemisinin, with or without 30 µM tadalafil. Right: gamete egress observed by IFAs. Samples were co-stained with mouse anti-glycophorin A (GPA, red) and rabbit anti-Pfg27 (green) IgG. DIC: Differential Interference Contrast. Arrow: mature GIE with an intact erythrocyte membrane, arrowhead: egressed gamete. Scale bars: 5 μm. Circles indicate the number of independent experiments and error bars show the SEM. Statistical significance is determined by one-way ANOVA with Dunnet correction (b, c, d, g, i) or with Sidak correction (f, h, j) for multiple comparisons or by a Mann Whitney test (e).

### cAMP-mediated reactivation of NPP increases uptake of artemisinin

Next, we analyzed whether the uptake of artemisinin derivatives is also regulated by cAMP-signaling. In early GIE, we observed that both pharmacological and genetic inhibition of PKA activity decreased the proportion of Fluo-DHA-positive cells (Fig. 4g and Fig. 4h), suggesting that PKA activity facilitates Fluo-DHA uptake in immature GIE. Thus, we hypothesized that a reactivation of NPP activity in mature GIE may increase their susceptibility to these drugs. First, we addressed whether activating cAMP pathway may enhance drug uptake by stage V GIE. Pre-incubation of mature GIE with the PDE inhibitors sildenafil and tadalafil strongly increased the proportion of Fluo-DHA-positive cells, reaching the level observed in early GIE (Fig. 4i and Supp. Fig. S6). To investigate the gametocytocidal effect of combining PDE inhibitors and artemisinin, we performed a gamete egress assay based on the ability of functionally viable mature gametocytes to undergo a temperature-dependent release of gametes from their host erythrocytes (*33*). We observed that the incubation of mature GIE with a combination of 5 µM artemisinin and 30 µM tadalafil drastically reduced the proportion of egressed gametes compared to tadalafil or artemisinin alone, indicating that phosphodiesterase inhibitors potentiate the effect of artemisinin in mature gametocytes (Fig. 4h). Therefore, these results show that cAMP-mediated reactivation of NPP in mature stages enhances uptake of artemisinin, and importantly this mechanism can increase mature gametocyte sensitivity to this antimalarial.

## DISCUSSION

It has been known for long that *P. falciparum* asexual stages modify their host erythrocyte to make them more permeable to supplementary nutrient uptake from the plasma and for removal of toxic waste by activating NPP in the erythrocyte membrane. In this study we unravel how these NPP are regulated during sexual parasite development and we propose that interfering with these pathways may facilitate artemisinin uptake by mature gametocytes. Our results raise a new model (Figure 5) where NPP activity in immature GIE depends on the PKA-mediated phosphorylation of one or several proteins directly or indirectly involved in channel activity. According to this model, NPP contribute to the uptake of artemisinin and derivatives by immature GIE. Then in mature GIE, PfPDEδ is highly expressed and degrades cAMP (*26*), leading to a drop of cAMP level that may reduce the phosphorylation of these proteins and consequently decrease NPP activity and drug uptake. Importantly, pharmacological inhibition of PfPDEδ can reactivate NPP activity and restore uptake of artemisinin derivatives by mature gametocytes, leading to an increased susceptibility to these drugs. Altogether these results show for the first time that *P. falciparum* gametocytes regulate infected erythrocyte permeability and that this permeability may facilitate uptake of some antimalarials.

**Figure 5.**
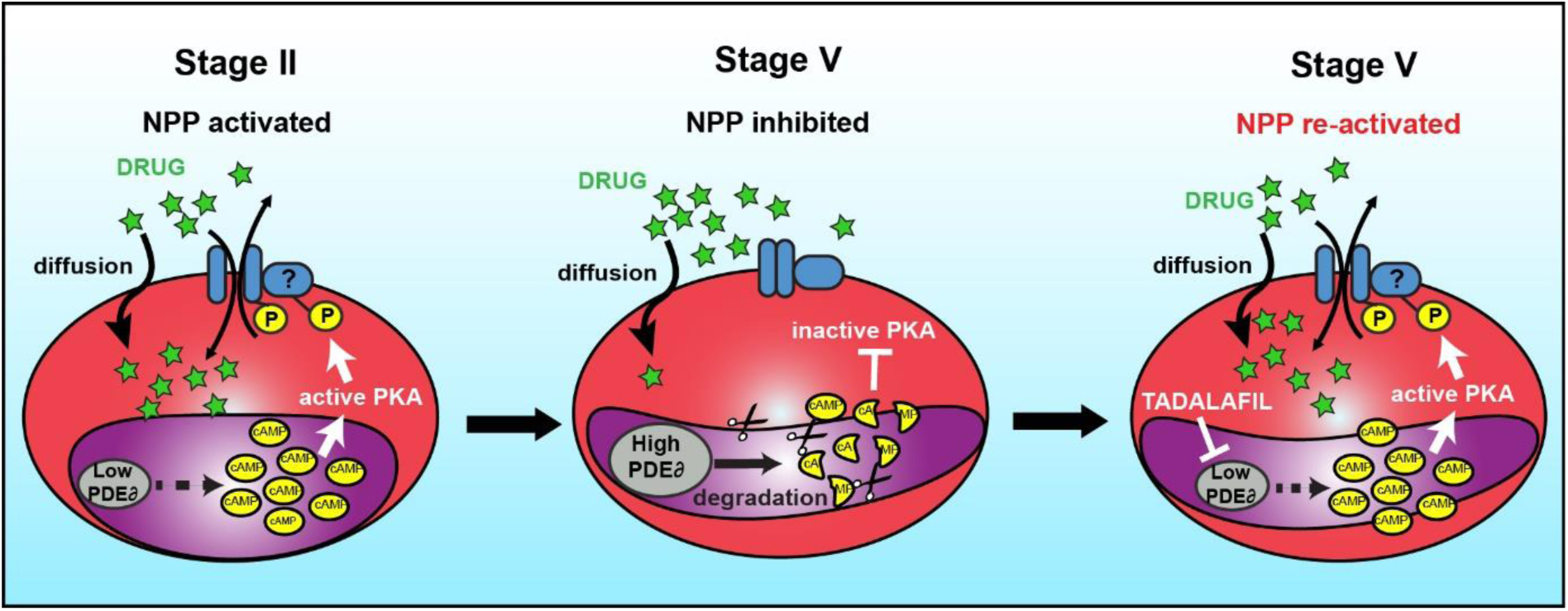
Model for cAMP-mediated regulation of NPP activity. Left: In early GIE, PfPDEδ expression is low, resulting in high cAMP levels and activation of PKA. PKA phosphorylates one or several proteins directly or indirectly involved in NPP activity that contributes to the uptake of drugs. Middle: In mature GIE, PfPDEδ is highly expressed and degrades cAMP, leading to a decrease in PKA phosphorylation and NPP activity. Right: In mature GIE, inhibition of PfPDEδ by tadalafil results in increased levels of cAMP that reactivate PKA and NPP, thereby restoring uptake of drugs.

Our results showing that NPP are still active in immature GIE contradict the paradigm that NPP activity is abolished in sexual stages. This dogma was based on a single study showing refractoriness of GIE to lysis upon short exposition to isosmotic sorbitol solution (*17*). In contrast, our data report that GIE were lysed after longer exposition to sorbitol, indicating the presence, although slower than in trophozoites, of NPP activity in immature GIE. In support to existence of NPP in sexual stages, recent RNA sequencing analyzes highlighted that sexually-committed parasites show higher transcription of *RhopH* and *clag* genes (*34*), whose products are involved in NPP activity (*12–14*). These observations suggest that members of the RhopH complex synthesized in sexually-committed parasites may be discharged upon egress onto the membrane of the erythrocyte targeted for invasion, within which a gametocyte will develop. This hypothesis would be consistent with the fact that gametocytes should absorb nutrients from the extracellular medium, such as panthotenate or isoleucine, for which NPP is the major route (*21*). In addition, NPP have been proposed to play a role in exporting amino acids liberated by the digestion of hemoglobin from the infected erythrocyte, thereby protecting the cell against the osmotic challenge posed by elevated intracellular amino acid levels (*20*). Thus, the slowdown of NPP activity in mature GIE may result from the completion of hemoglobin digestion in these stages (*35*). Absence of channel activity in mature stages may allow the infected erythrocyte to decrease its osmotic fragility and thus to persist longer in the blood circulation. This process appears to be tightly regulated by cAMP signaling pathway, as previously observed for asexual stages (*16*).

These results are in accordance with the detection of CLAG3/RhopH1, one of the major component of NPP, in a global phospho-proteomic analysis of *P. falciparum* parasites *(36).* As part of the cAMP signaling cascade, the PfPDEδ enzyme, whose expression increases in mature gametocytes (*26*), plays an instrumental role in the NPP decline. Interestingly, this enzyme has also been reported to govern the switch in GIE deformability that enables mature gametocytes to circulate several days in the bloodstream and avoid the clearance by the spleen (*26, 37, 38*). Therefore, by decreasing GIE osmotic fragility and increasing GIE deformability, PfPDEδ appears to be the key regulator of mature gametocytes persistence in the blood circulation. Importantly, our present work suggests that targeting this enzyme with the FDA-approved drug tadalafil may enhance artemisinin uptake by mature gametocytes, thereby decreasing their refractoriness to this antimalarial. As PDE inhibitors also render mature GIE rigid and hence may promote their clearance by the spleen (*26*), they represent novel drug leads potentially capable of blocking malaria transmission by impacting on both gametocytes circulation and susceptibility to artemisinin derivatives.

## MATERIAL AND METHODS

### Parasite culturing and gametocyte production

The *P. falciparum* NF54 strain, the B10 clone and the transgenic lines pHL*pfpkar*, PfPDEδ^−^, NF54-cg6-pfs16-CBG99 and NF54-pfs47-pfs16-GFP have been described elsewhere (*16, 23, 24, 27, 39*). Parasites were cultivated *in vitro* under standard conditions using RPMI 1640 medium supplemented with 10% heat-inactivated human serum and human erythrocytes at a 5% hematocrit. To obtain synchronous asexual stages, parasites were synchronized by the isolation of schizonts by magnetic isolation using a MACS depletion column (Miltenyi Biotec) in conjunction with a magnetic separator, and placed back into culture. After invasion of merozoites, a second magnetic isolation was used for the selection of ring-stage parasites to obtain a tighter window of synchronization. Synchronous production of specific gametocytes stages was achieved by treating synchronized cultures at the ring stage (10-15% parasitemia, day 0) with 50 mM N-acetylglucosamine (NAG) for 5 days to eliminate asexual parasites. Gametocyte preparations were enriched in different experiments by magnetic isolation. Stage I GIE were collected at day 1 post NAG treatment, stage II GIE were collected at days 2 and 3, stage III GIE were collected at days 4 and 5, stage IV GIE were collected at days 6 and 7, and stage V GIE were collected at day 8 onwards.

### Isosmotic lysis

500 µl of *P. falciparum* cultures (containing gametocytes or asexual stages) with a parasitemia >0.5% were washed once with RPMI and incubated in 1.5ml tubes for 60 min at 37°C in 500µl isosmotic solution containing either 300 mM sorbitol, 300 mM Alanine or 150 mM PhTMA^+^ supplemented with 10 mM Hepes, 5 mM glucose and with a pH adjusted to 7.4. For each experiment, 5 tubes were prepared including one tube containing 100 µM NPPB, Furosemide, Ro5-4864, or PK11195 (all purchased from Sigma-Aldrich). At each sampling time, one tube was spinned and smears were prepared and stained with Giemsa R solution (RAL diagnostics). Parasitemia was estimated for each point by counting infected cells out of at least 4000 erythrocytes, and lysis percentage was calculated using the formula: % lysis (t)= [1-(parasitemia (t)/parasitemia (t0))]*100

The percentage of lysis of stage I GIE was determined using the NF54-pfs47-pfs16-GFP line by fluorescence microscopy at X40 magnification on a Leica DMi8 microscope. To address the role of cAMP-signaling in isosmotic lysis, stage II GIE (day 2 and 3 post NAG treatment) were pre-incubated 30 minutes with 100 µM H89, 10 µM KT5720 or 100 µM 8Br-cAMP, and mature GIE (day 8-11 post NAG treatment) were pre-incubated 30 minutes with tadalafil or sildenafil at different concentrations (from 10 µM to 100 µM). All inhibitors were purchased from Sigma-Aldrich or Euromedex.

### Patch-clamp

Patch clamp experiments were performed at room temperature using the whole cell configuration. Pipettes were pulled using a DMZ Universal Puller (Zeitz Instruments, Germany) from borosilicate glass capillaries (GC150F-10, Harvard apparatus, UK) to obtain a tip resistance of 10-15 MΩ. Pipette solution was: 145 mM KCl, 1.85 mM CaCl_2_, 1.2 mM MgCl_2_, 5 mM EGTA, 10 mM Hepes, 10 mM, pH 7.2. Bath solution was: 145 mM NaCl, 5 mM KCl, 1 mM CaCl_2_, 1 mM MgCl_2_, 10 mM Hepes, 10 mM glucose, pH 7.4. Seal resistance were 4-20 GΩ, and whole cell configuration was obtained by a brief electrical pulse (zap) and was assessed by development of small capacitance transient currents and reduction of access resistance. Whole cell current were recorded using either an Axopatch 200B (Molecular Devices, USA) or a RK400 (Biologic, France) amplifier, with voltage command protocols and current analysis done with pclamp10 suite software (Molecular Devices, USA) or WinWCP4.7 software (J. Dempster, Strathclyde University, UK), respectively. Currents were elicited by generating a series of membrane potentials from +100 to −100 mV in −10 mV steps for 500 ms, from a holding potential of 0 mV. For stage I GIE, the gametocyte fluorescent NF54-pfs47-pfs16-GFP line was used to easily distinguish by microscopy between asexual and early sexual stages.

### Fluorescence microscopy

GIE were incubated 2 hours with Fluo-DHA or DMSO 0.1 % at 37 °C. Cells were stained with Hoechst 33342 (1/20,000, Thermo Fisher Scientific) for 10 min at 37 °C. After one wash with Phosphate-Buffered-Saline (PBS) 1X, cells were fixed for 10 min at room temperature with 1 % paraformaldehyde (PFA) (Electron Microscopy Science) and 0.25 % glutaraldehyde (Sigma-Aldrich) in PBS 1X. After three washes with PBS 1X, infected cells were observed on a glass slide at X100 magnification using a Leica DMi8 microscope.

### Quantification of Fluo-DHA uptake assay

The synthesis of the Fluo-DHA probe was described in Sissoko *et al* (*31*). Quantification of Fluo-DHA uptake was performed using flow cytometry. GIE were pre-incubated or not with 100 µM NPPB or with 100 µM Furosemide for 30 minutes and then incubated with 1 µM of Fluo-DHA for 2 hours at 37°C, 5% CO2 and 5% O2. To address the role of cAMP-signaling in Fluo-DHA uptake, early GIE (day 2 post NAG treatment) were pre-incubated 30 minutes with 100 µM H89, 10 µM KT5720 or 100 µM 8Br-cAMP, and mature GIE (day 8-11 post NAG treatment) were pre-incubated 30 minutes with tadalafil or sildenafil at different concentrations (from 10 µM to 100 µM). To assess the specificity of Fluo-DHA uptake, early GIE were pre-incubated or not with 20 µM DHA for 2 hours and then incubated with Fluo-DHA (1 µM) for 2 hours. DHA alone (20 µM) and HPA-NBD (1 µM) were used as control. Twenty minutes before the end of incubation, GIE were stained with Hoechst 33342 (1/10,000). Cells were then washed with PBS 1 X and fixed for 10 min at room temperature with PBS, 1 % PFA and 0.25 % Glutaraldehyde. The percentage of GFP positive cells were quantified using Fortessa (BD Biosciences) cytometer.

### Gametocyte survival assay

To calculate the IC_50_ for Fluo-DHA on early gametocytes, 2.10^5^ MACS-purified early GIE (day 2 post-NAG treatment) from the NF54-cg6-Pfs16-CBG99 line were incubated with serial dilutions of Fluo-DHA for 3 hours, then GIE were washed and incubated with complete medium without drugs for 48 hours. To perform viability assays with drugs and inhibitors, early GIE were pre-incubated at 37°C for 30 minutes with 100 µM NPPB or 0.1 % DMSO in complete medium and then incubated with the same inhibitors supplemented with 700 nM artemisinin (Sigma-Aldrich), 150 nM Fluo-DHA or 0.1 % DMSO for 3 hours. GIE were then washed and incubated with complete medium without inhibitors and drugs for 48 hours. Cell viability was evaluated by adding a non lysing formulation of 0.5 mM D-Luciferin substrate (*23*) (Sigma-Aldrich) and measuring luciferase activity for 1 second on a plate Reader Infinite 200 PRO (Tecan®). All experiments were performed in triplicate on 96 well plates.

### Gamete egress assay

NF54 cultures containing 2% to 5% stage V gametocytes were incubated for 24 h at 37°C with 5 µM artemisinin supplemented or not with 30 µM tadalafil. Controls comprising gametocytes exposed to the same final concentration of DMSO (0.1%) were processed in parallel. After the incubation, 100 µl samples from each condition were pelleted at 1,000 g for 1 min and rapidly resuspended in 50 µl human serum at room temperature for 30 minutes during which gametogenesis took place. Samples were then pelleted at 1,000 g for 1 min, smeared on glass slides, methanol-fixed and co-stained with mouse anti-glycophorin A (GPA, 1/1,000, Santa Cruz Biotechnology) and rabbit anti-Pfg27 (*40*) (1/2,000) followed by anti-rabbit Alexa 488- and anti-mouse Alexa 594-conjugated IgG (1/2,000, Life technologies). Samples were observed at 100X magnification using a Leica DMi8. At least 100 GIE were analyzed for each sample. Percent gamete egress was determined by calculating the % of GPA-negative round gametes in the total GIE population detected by Pfg27 staining.

## Acknowledgements

The authors thank D. Baker (LSHTM) for providing the PfPDEδ^−^ line. The authors acknowledge the Flow Cytometry core facility CYBIO of the Institut Cochin for technical help. This study was supported by grants from Laboratory of Excellence GR-Ex, reference ANR-11-LABX-0051. The labex GR-Ex is funded by the IdEx program “Investissements d’avenir” of the French National Research Agency, reference ANR-18-IDEX-0001. CL, DB, FD, AM, MEN, GN, and LB acknowledge the financial support from the Cnrs, Inserm and the Fondation pour la Recherche Médicale (“Equipe FRM” grant EQ20170336722).

## Author contributions

G.B., S.E. and C.L. conceived the project. G.B., D.B., J.C., R.D., S.E. and C.L. designed and interpreted the experiments. G.B., D.B., F.D., A.M., A.S., M-E.N., G.N., L.B., N.K., D.R. and C.L. performed the experiments. S.H., G.S., P.A., R.M.M., J-J.L-R. and R.D. contributed resources or data. C.L. wrote the article, with major input from G.B., J.C., R.D. and S.E.

