## Supplementam material for "*Plasmodium falciparum* sexual parasites regulate infected erythrocyte permeability"

**SUPPLEMENTAL FIGURES AND LEGENDS**

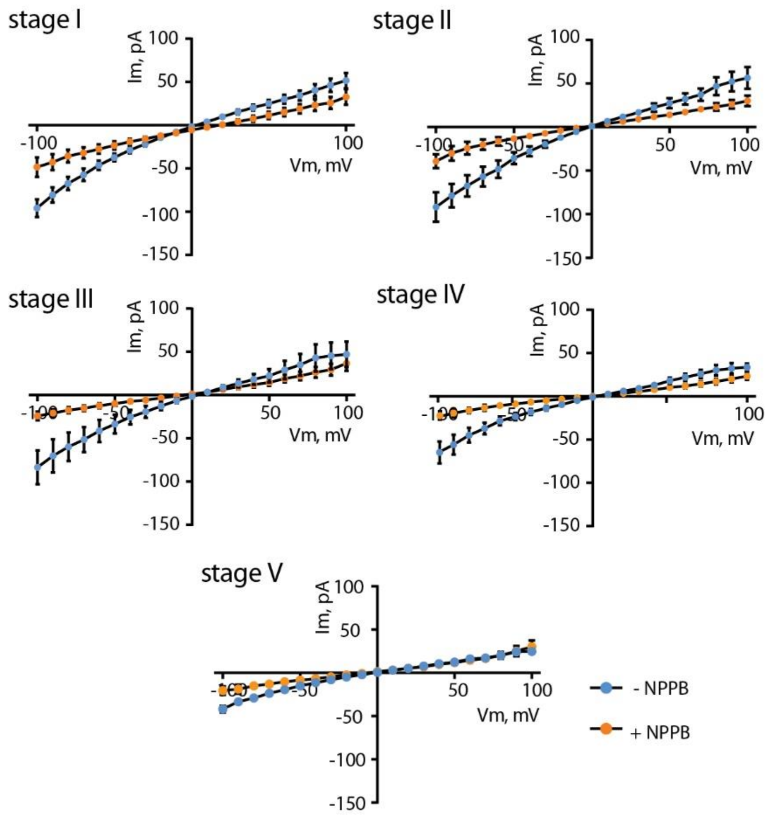


**Supplemental Figure S1**

I-V plots (B) from patch experiments on GIE from stage I to stage V with (orange) or without (blue) 100 µM NPPB.

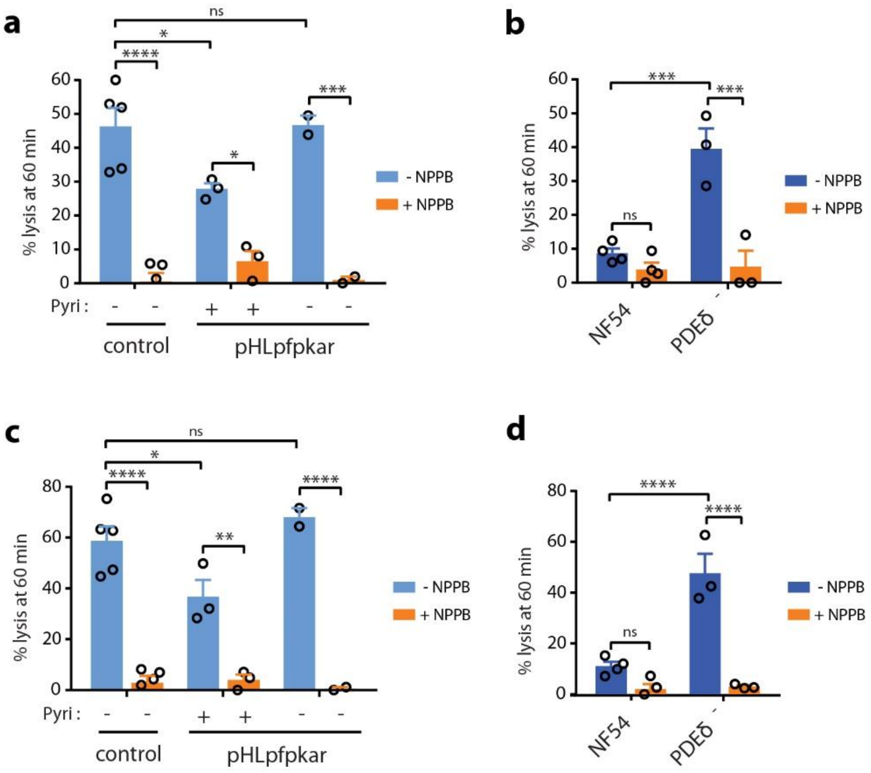


**Supplemental Figure S2**

**a, c.** Alanine-induced (a) and PhTMA$^+$-induced (c) isosmotic lysis of stage II GIE from the NF54 isolate (Control) and the transgenic pHL*pfpkar* line, cultivated with and without pyrimethamine (Pyri).

**b, d.** Alanine-induced (b) and PhTMA$^+$-induced (d) isosmotic lysis of stage V GIE from the NF54 isolate and the transgenic line PDEδ$^-$. All experiments were performed in presence or absence of 100 µM NPPB. Circles indicate the number of independent experiments, error bars show the SEM and statistical significance is determined by one-way ANOVA with Sidak correction for multiple comparisons.

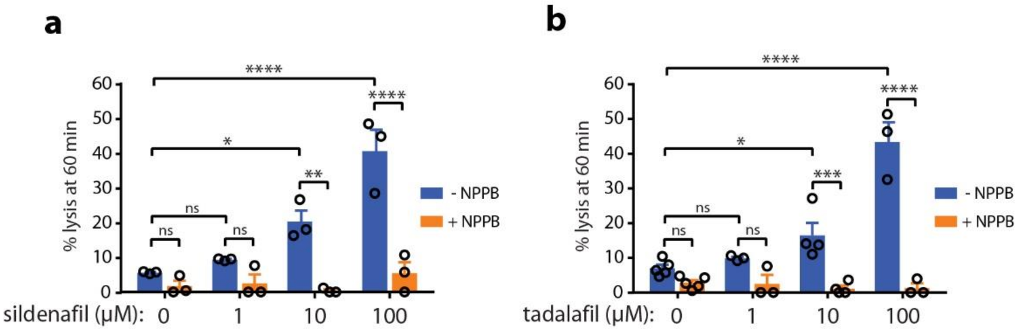


**Supplemental Figure S3.**

Sorbitol-induced isosmotic lysis of stage V GIE with 1 µM, 10 µM or 100 µM sildenafil (a) or 100 with 1 µM, 10 µM or 100 µM tadalafil (b). All experiments were performed in presence or absence of 100 µM NPPB. Circles indicate the number of independent experiments, error bars show the SEM and statistical significance is determined by one-way ANOVA with Sidak correction for multiple comparisons.

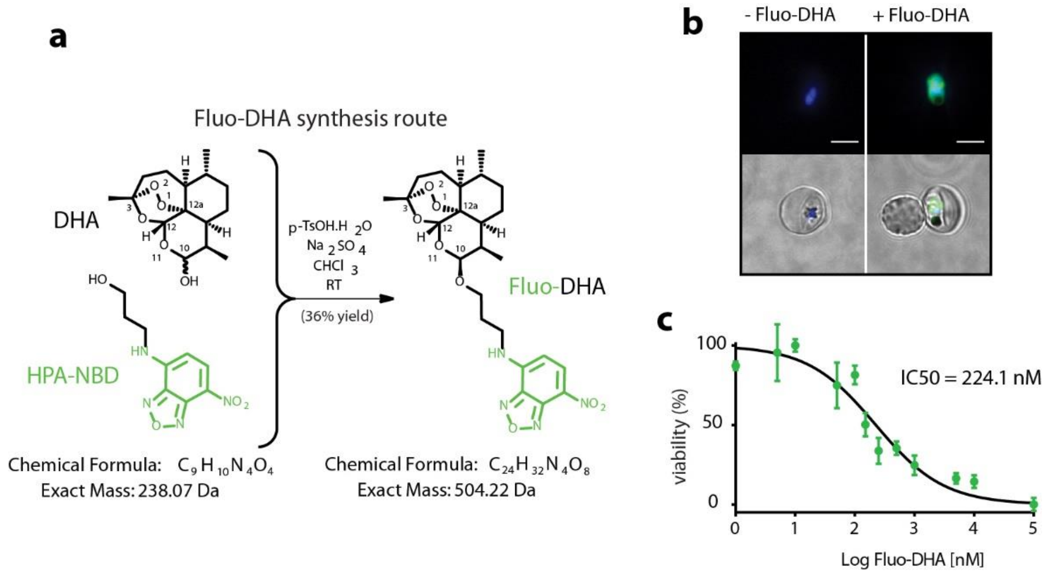


**Supplemental Figure S4.**

**a.** Schematic of Fluo-DHA synthesis from dihydroartemisinin (DHA) and the fluophore HPA-NBD. **b.** Fluorescence microscopy imaging of paraformaldehyde-fixed early GIE showing Fluo-DHA uptake (green). DNA is stained with Hoechst 33342 (blue). Scale bars: 5 µm. **c.** Dose-response curve and $IC_{50}$ value for Fluo-DHA in early gametocytes from the NF54-cg6-pfs16-CBG99 line. Viability (luciferase activity) was determined 48 hours after a 3-hour incubation with serial dilutions of Fluo-DHA. Error bars show the SEM.

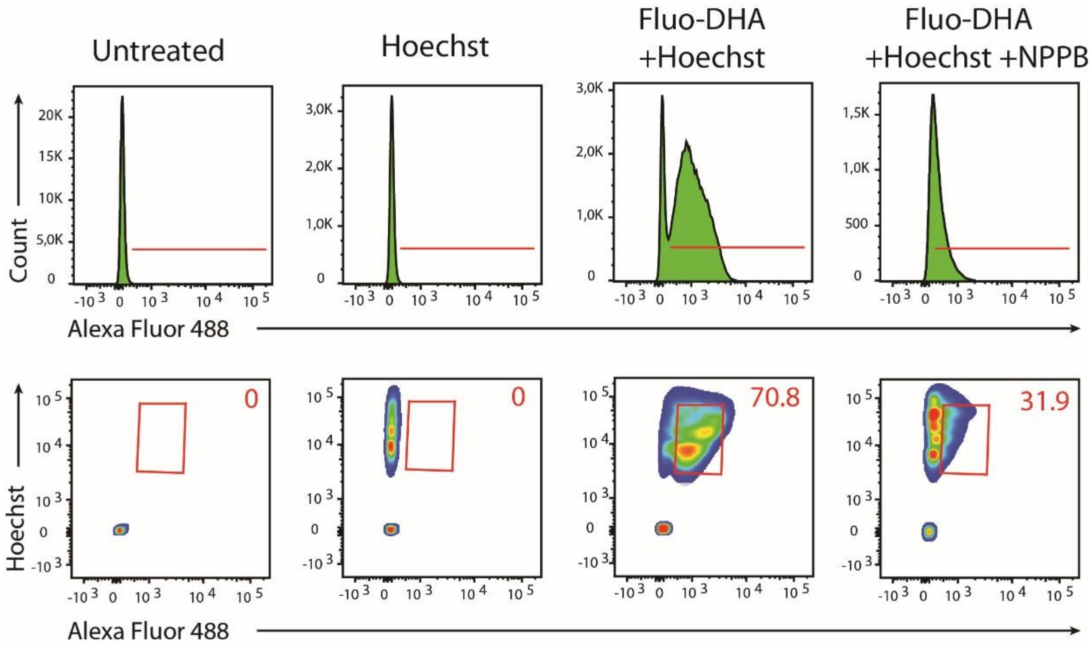


**Supplemental Figure S5.**

Scatter plots showing the gating strategy for Fluo-DHA uptake. Early GIE were pre-incubated or not with 100 µM NPPB for 30 minutes and then incubated with 1 µM of Fluo-DHA for 2 hours. Twenty minutes before the end of incubation, GIE were stained with Hoechst 33342. Percentages of Fluo-DHA uptake are shown.

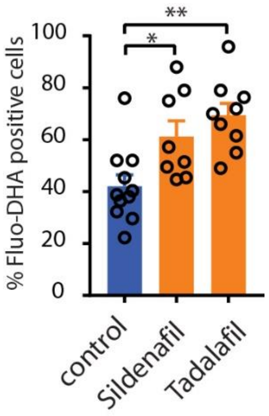


**Supplemental Figure S6.**

Fluo-DHA uptake in stage V GIE upon 100 µM tadalafil or 100 µM sildenafil incubation. Circles indicate the number of independent experiments, error bars show the SEM and statistical significance is determined by one-way ANOVA with Dunnett correction for multiple comparisons.
